# Increased diffusion in livers with advanced fibrosis: pre-clinical and clinical observations with diffusion MRI

**DOI:** 10.64898/2026.03.30.715426

**Authors:** Fan-Yi Xu, Yì Xiáng J. Wáng

## Abstract

Despite the increased water content in fibrotic livers, numerous studies reported a decrease in ADC (apparent diffusion coefficient) in liver fibrosis. We argue that the ADC decrease in fibrotic livers is due to the ‘T2 shine-through’ of ADC, as the longer T2 in liver fibrosis leads to less signal decay between the low and high *b*-value images. The metric slow diffusion coefficient (SDC) was proposed to mitigate the difficulties associated with this ‘T2 shine-through’ of ADC. This study calculated ADC and SDC of one rat study with liver fibrosis induced by biliary duct ligation (BDL), and three sets of human liver fibrosis data. To tease out the menopausal effect on SDC, only the results of men’s livers were analysed for the human datasets. The rat study showed, liver ADC decreased stepwise (in weeks after BDL procedure) following fibrosis induction, SDC increased stepwise. In human studies, all three datasets consistently showed advanced fibrosis had an ADC lower than that of earlier stage fibrosis; advanced fibrosis had a SDC higher than that of earlier stage fibrosis. When each liver SDC datum was normalized by the mean value of the controls without fibrosis, and the three human datasets were summed together, stage-1 liver fibrosis had a normalized SDC value lower than that of the controls, and there was a stepwise increase of SDC value from stage-1 liver fibrosis to stage-4 liver fibrosis. It is known that liver fibrosis is associated with lower perfusion, higher iron/susceptibility, and higher water content, and these three factors all contribute to the lower ADC measure. Higher iron/susceptibility lowers SDC measure, whereas higher water content elevates SDC measure. It is likely that for early-stage fibrosis, the net effect of susceptibility and water leads to a lower SDC, while for advanced fibrosis, the net effect leads to a higher SDC.

---

Patients with advanced liver fibrosis show an abnormal regulation of extracellular fluid volume, resulting in the accumulation of sodium and water retention, ascites, oedema and pleural effusion. Ascites commonly occurs in diseases causing sinusoidal portal hypertension such as liver cirrhosis (i.e., stage 4 liver fibrosis) [1, 2]. Schober *et al*. [3] reported that, compared with control subjects, extracellular water (ECW), total body water (TBW), intracellular water, plasma volume all increased in patients with cirrhosis. With a 1.5 T scanner, Mesropyan *et al*. [4] reported that, compared with those of healthy controls, cirrhotic livers with portal hypertension had longer T2 (53.72 ± 7.56 ms vs. 48.58 ± 8.41 ms) and higher extracellular volume fraction values (45 ± 18.55% vs 26.14 ± 2.31%). Higher liver water extent can be seen in earlier stages of liver fibrosis. In a human study, Guimaraes *et al*. [5] described that liver T2 value (1.5 T) was: control 65.4±2.9 ms; mild fibrosis (Ishak grade 1–2) 66.7±1.9 ms; moderate fibrosis (Ishak grade 3–4) 71.6±1.7 ms; severe fibrosis (Ishak grade 5–6) 72.4±1.4 ms. In an animal study where diethylnitrosamine (DEN) was used to include fibrosis in rats, Guimaraes *et al*. [5] described a monotonic increase in T2 (4.7T) between each rat subgroup, with phosphate buffer solution rats 25.2±0.8 ms, DEN 5-week exposure rats 31.1±1.5 ms, and DEN 8-week exposure rats 49.4±0.4 ms. Liver fibrosis is associated with elevated ECW. Applying bioimpedance analysis for viral hepatitis related liver diseases, Nishikawa *et al.* [6] reported that the median of ECW/TBW [ratio of ECW to TBW] was: 0.381 in fibrosis stage 0-1, 0.384 in fibrosis stage 3, 0.389 in Child-Pugh A, and 0.395 in Child-Pugh B or C. Kishino *et al.* [7] reported ECW/TBW ratio was positively correlated with Fibrosis-4 Index and APRI (Aspartate Aminotransferase-to-platelet ratio index) and negatively correlated with albumin value and prothrombin time.

Despite the increased water content in fibrotic livers and particularly in cirrhotic livers, it is paradoxical that numerous studies reported a decrease in ADC (apparent diffusion coefficient) in liver fibrosis and particularly in liver cirrhosis [8–12]. Girometti *et al.* [9] described that liver ADC was significantly lower in cirrhotic livers than in controls (1.11±0.16 vs. 1.54±0.12 ×10^-3^ mm^2^/s). In the study reported by Kahraman *et al*. [10], patients with chronic liver disease were classified as group 1, group 2, and group 3 according to the absence of ascites, the presence of minimal ascites, and the presence of massive ascites, respectively. In the control group, liver ADC value was 1.04 ± 0.11×10^-3^ mm^2^/s. In patients with chronic liver disease, liver ADC was 0.90 ± 0.10 × 10^−3^ mm^2^/s for group 1, 0.87 ± 0.09 × 10^−3^ mm^2^/s for group 2, and 0.79 ± 0.09 × 10^−3^ mm^2^/s for group 3. Subbiah *et al*. [11] also reported that End-Stage Liver Disease (MELD) score was negatively correlated with liver ADC.

Recently, the ‘T2 shine-through’ effect of ADC has been described (also see the discussion section) [13–16]. The application of diffusion gradients leads to the tissue demonstrating shorter ‘apparent’ (or ‘measured’) T2 relative to the T2 measured when *b*= 0 s/mm^2^. In a study of breast cancer tissues (3.0T), Egnell *et al* [17] reported that the ‘apparent’ T2 value at *b* = 50 s/mm^2^ was around 8% lower than at *b* = 0 s/mm^2^ (68.7 ms vs 74.5 ms). This phenomenon can be more apparent when the actual T2 of a tissue is short (such as < 70 ms), and this can explain many of the puzzling observations. For example, solid cartilage has a high ADC (1.5 ×10^-3^ mm^2^/s) due to its very short T2 (37 ms, 3.0T) allowing a fast signal decay between the low *b*-value image and the high *b*-value image [14, 16]. The spleen (with a T2 of around 60 ms at 3.0 T, ADC of only around 0.8 ×10^-3^ mm^2^/s) and parotid gland Warthin’ tumors (with a T2 of around 80 ms at 3.0 T) have low ADCs despite having rich blood perfusion [16, 18]. This ‘T2 shine-through’ effect of ADC is observed regardless of whether *b*=0 mm^2^/s image is included for ADC calculation or not [13, 14]. The new DWI metric slow diffusion coefficient (SDC) was proposed to mitigate the difficulties associated with this ‘T2 shine-through’ of ADC [19]. In its basic form, SDC is derived from the subtraction of a high *b*-value DWI image (e.g., *b*=400 s/mm^2^) and a higher *b*-value DWI (e.g., *b*=600 s/mm^2^). It is known that the spleen has a lower ADC than the liver, hepatocellular carcinoma (HCC) has a lower ADC than liver parenchyma, liver hemangiomas have a lower ADC than simple liver cysts. With SDC analysis, the spleen has a faster diffusion than the liver, HCC has faster diffusion than liver parenchyma, liver hemangiomas have a faster diffusion than simple liver cysts [19]. The liver and spleen have a similar amount of blood perfusion, the spleen is waterier than the liver. HCCs are mostly associated with increased blood supply and increased proportion of arterial blood supply and with edema. It is more reasonable with SDC results that spleen and HCC have a faster diffusion than liver parenchyma. Due to the ‘flushing’ of blood flow inside the hemangioma, it is also more reasonable with SDC results that the diffusion of hemangioma liquid is faster than the more ‘static’ liquid of the cysts.

In this study, we demonstrate that while advanced liver fibrosis is associated with a lower ADC, advanced liver fibrosis is associated with an elevated SDC. The SDC result is consistent with the known pathophysiology that advanced liver fibrosis is associated with higher water content and more so with ECW content and thus faster tissue diffusion.

## Materials and Methods

This study re-utilized one previously published animal liver IVIM study dataset [20], two published human liver IVIM study datasets [21, 22], and one newly acquired human liver IVIM study dataset. All MRI data acquisitions were approved by the local institutional ethical committees, and the informed consent was obtained for all the human subjects. The animal experiment was approved by the local animal research committee. In a recent healthy volunteer study, we noted that SDC measure is highly sensitive to the iron-related susceptibility effect. Healthy women have a higher liver SDC measure than that of healthy men, and post-menopausal women have a lower liver SDC measure than that of pre-menopausal women. Note that men’s liver has a higher liver iron level than that of women, and post-menopausal women have a higher liver iron level than that of pre-menopausal women. On the other hand, men’s liver SDC measures remain stable across age groups [23]. Thus, in the current study, only the results of men’s livers were analysed and presented. Due to the lack of sufficient patient number for each fibrosis stage, in this study, stage 3 and stage 4 in human dataset 1 and dataset 2, and MRI visible cirrhosis were termed ‘advanced stage’ liver fibrosis, and stage 1 and stage 2 in human dataset 1 and dataset 2, and liver fibrosis but without MRI visible cirrhosis were termed ‘early-stage’ liver fibrosis.

### Rat fibrosis model induced by biliary duct ligation (BDL)

The study for this animal data has been described earlier [20]. For the biliary duct ligation (BDL) model, Sprague-Dawley rats (weight: 300 ± 20g) were anesthetized, and an upper abdominal incision made, the lower segment of common bile duct ligated. Recanalization was performed with rats having BDL for 7 days. Via a laparotomy, the ligated distal end of the common bile duct was dissected, and an anastomosis was established between the common bile duct and the jejunum, which was sectioned 5 cm from the duodenojejunal angle. MR images were acquired with a clinical 3.0T magnet (Ingenia, Philips Healthcare, Best, Netherlands) using a 4-channel small animal coil. The IVIM type of diffusion scan was based on a single-shot spin-echo type echo-planar sequence, and the parameters were as follows: TR/TE = 2000/55 ms; FOV=50×50 mm, slice thickness = 3 mm, number of slices = 9, matrix = 64×63. SPIR technique (Spectral Pre-saturation with Inversion-Recovery) was used for fat suppression. DWI images with *b*-values of 0, 800, 1000 s/mm^2^ were used in the current study. NEX was 3 and 4 respectively for images of *b*=800 and *b*=1000, while NEX was 1 for images of *b*=0 s/mm^2^. In addition to 9 control rats, the rat number was 5, 6, 5, and 3 respectively, for the timepoints of 1 week, weeks, 3 weeks, 4 weeks post-BDL surgery. Recanalization was performed with rats having BDL for 7 days. MRI for recanalization rats was performed with 1 week (n=5) or 2 weeks(n=6) after the recanalization procedure. Each animal was MRI scanned once. Animals were sacrificed within 4 hours after MRI, and liver specimens were taken for the histological assessment of liver fibrosis

### Human viral B hepatitis related fibrosis data 1

The human dataset 1 has been described earlier [21]. The IVIM type of diffusion scan was based on a single-shot spin-echo type echo-planar sequence using a 1.5-T magnet (Achieva, Philips Healthcare, Best, Netherlands). SPIR technique was used for fat suppression. Respiratory-gating was applied. The TR was 1600 ms and the TE was 63 ms, with one TR per respiratory cycle (hereby TR refers to the time from radiofrequency pulse to echo signal acquisition). Other parameters included slice thickness =7 mm and inter-slice gap 1mm, matrix= 124×97, FOV =375 mm×302 mm, NEX=2, number of slices =6. DWI images with *b*-values of 0, 400, 600 s/mm^2^ were used in the current study. In total there were 13 healthy volunteers (mean age: 23.85 years, range: 20 to 31 years), 7 early-stage fibrosis patients (mean age: 42.4 years, range: 23 to 65 years), and 4 advanced fibrosis patients (mean age: 51.5 years, range: 32 to 60 years).

### Human viral B hepatitis related fibrosis data-2

The human dataset 2 was newly acquired for the current study. Using a 1.5-T magnet (uMR, United Imaging Healthcare, Shanghai, China), the IVIM type of diffusion scan was based on a single-shot spin-echo type echo-planar sequence with free breathing [24]. SPIR technique was used for fat suppression. The TR was 1600 ms and the TE was 65 ms. Other parameters included slice thickness =7 mm and inter-slice gap 2 mm, matrix= 128×99, FOV =256 mm×204 mm, number of slices =12. DWI images with *b*-values of 0, 400,600 s/mm^2^ were used in the current study. NEX was 2 for images of *b*=400 and *b*=600, while NEX was 3 for images of *b*=0 s/mm^2^. In total there were 11 healthy volunteers (mean age: 37.2 years, range: 22 to 58 years), 11 early-stage fibrosis patients (mean age: 42 years, range: 15 to 65 years), and 9 advanced fibrosis patients (mean age: 52 years, range: 34 to 69 years).

### Human viral B hepatitis related fibrosis data-3

The human dataset 3 has been described earlier [22]. Liver IVIM imaging was performed with a 3.0-T magnet (Vida Magneton, Siemens Healthineers, Erlangen, Germany). The diffusion imaging was based on a single-shot spin-echo type echo-planar sequence with respiratory gating. The default spectral pre-saturation technique was used for fat suppression. The TR was 2500 ms and TE was 84 ms. Other parameters included slice thickness = 5 mm and inter-slice gap =1 mm, matrix = 128 × 128, FOV = 350 mm × 350 mm. DWI images with *b*-values of 0, 500, 800 s/mm^2^ were utilized in this study. NEX was 1 for *b*= 0 s/mm^2^ images, and 3 for *b*= 500, 800 s/mm^2^ images. For this dataset, the initial study aimed at diffusion MRI separation of FNH (focal nodular hyperplasia) and liver malignant tumors [22]. All patients have surgical histopathology. In the current analysis, the control group (n=6, mean age: 37.67 years, range: 22 to 45 years) was FNH patients who did not have a diffused liver disease background except the focal lesion; the patient group all had HCC and a viral hepatitis B related liver fibrosis background. We separated HCC patients into two groups: fibrosis group (n=28, mean age: 51.3 years, range: 40 to 59 years) and MRI visible cirrhosis group (n=9, mean age: 55.7 years, range 43 to 73 years). MRI visible cirrhosis patients had MRI visible cirrhosis sign(s) and/or had ascites plus apparent splenomegaly [25].

### Data analysis

ADC was calculated according to:

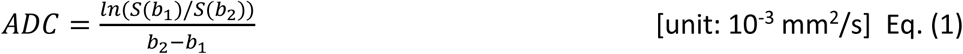

where *b_2_* and *b_1_* refer to high *b*-value and low *b*-value (*b*=0 s/mm^2^ in this study), respectively, S(*b_2_*) and S(*b_1_*) denote the image signal-intensity acquired at the high *b*-value and low *b*-value, respectively. *b_2_* was 1000 s/mm^2^ for the animal study, 600 s/mm^2^ for human study datasets 1 and 2, and 800 s/mm^2^ for human study dataset 3.

SDC was calculated according to [’9]:

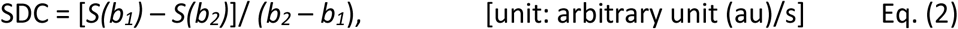

where *b_1_* and *b_2_* refer to a high *b*-value and a higher *b*-value respectively, where S(*b_1_*) and S(*b_2_*) denote the DW image signal-intensity acquired at the high *b*-value and the higher *b*-value respectively. *b_1_* was 800 for the animal study, 400 s/mm^2^ for human study datasets 1 and 2, and 500 s/mm^2^ for human study datasets 3. *b_1_* was 1000 for the animal study, 600 s/mm^2^ for human study datasets 1 and 2, and 800 s/mm^2^ for human study datasets 3. A higher SDC value indicates a more rapid signal decay between the two *b*-values, reflecting faster diffusion [19].

Image segmentation was performed using ITK-SNAP (http://www.itksnap.org) and data analysis was conducted with MATLAB (MathWorks, Natick, MA, USA). For ADC, free-hand ROIs (regions-of-interest) were manually placed on *b*=0 s/mm^2^ image to cover a large portion of liver parenchyma while avoiding large vessels (and avoiding focal lesion and focal lesion’s surrounding tissues for dataset-3) and then copied on to the high *b-*value images of this slice. To account for the potential inter-scan motion, the copied ROIs on the high *b-*value images were additionally manually adjusted. For SDC, free-hand ROIs were initially placed on the high *b-*value images, then copied on to the higher *b-*value images of this slice. The ROI drawing was conducted by a trained engineer graduate and checked by a specialist radiologist. For all analysis, the mean of all included slices’ measurements was regarded as the value of the examination, with the last step weighted by the percentage ROI area for each slice (i.e., assuming the sum pixel number of all ROIs for each subject being 100%, according to pixel number in each slice’s ROI, a percentage was assigned for each slice).

For statistical analysis, data were processed using GraphPad Prism (San Diego, CA, USA). Comparisons were performed using the Mann Whitney test or the Kruskal-Wallis test as appropriate. A P-value <0.05 was considered statistically significant.

## Results

The rat study showed, liver ADC decreased stepwise (measured in weeks after BDL procedure) following fibrosis induction, whereas SDC increased stepwise following BDL procedure. Following the recanalization, both the liver ADC and the liver SDC returned to the values of the controls (*Figure-1, table -1*).

**Figure-1.**
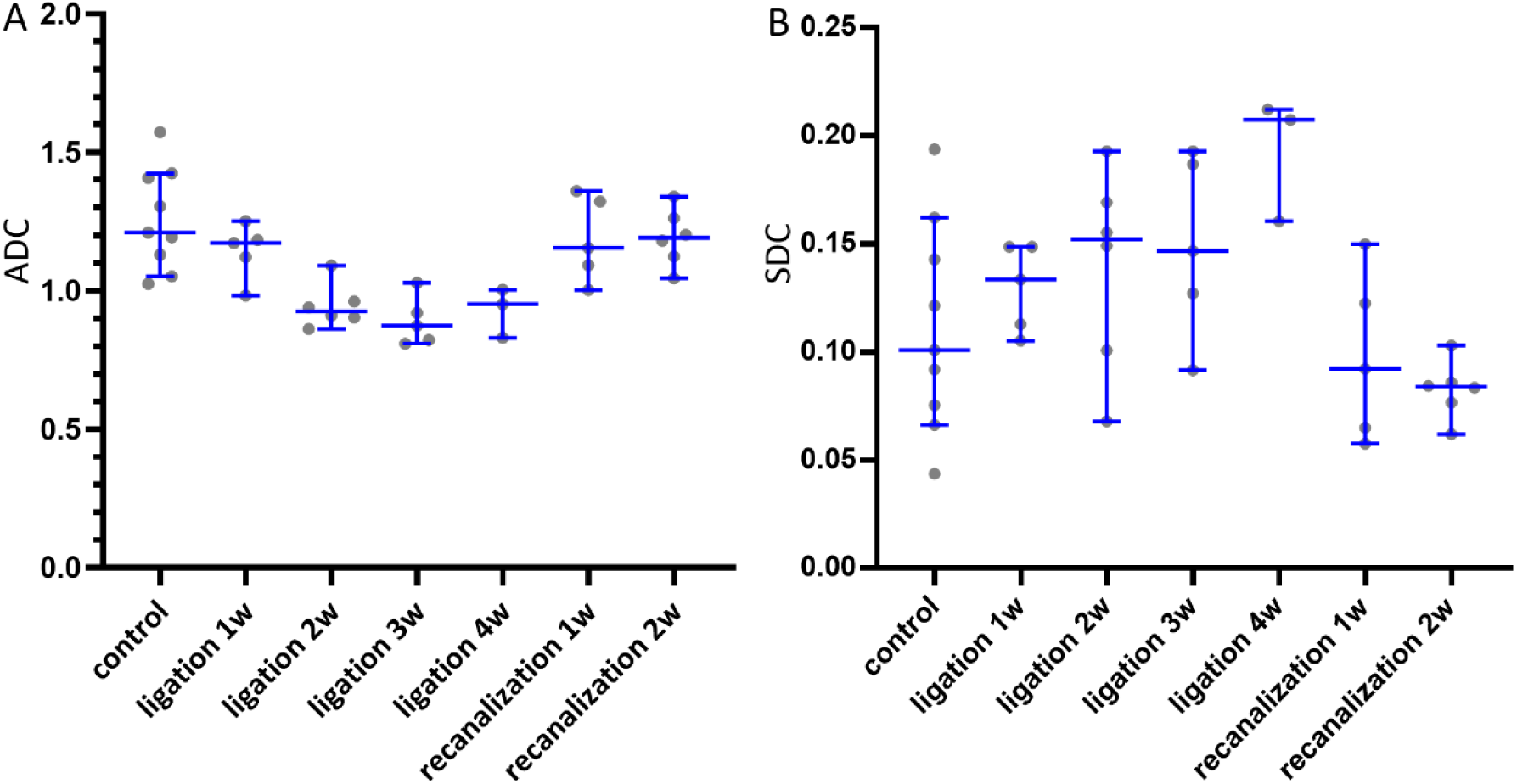
ADCb0b1000 and SDC_b800b1000_ changes following BDL procedure and following recanalization procedure. Liver ADC decreased stepwise (in weeks after BDL procedure) following fibrosis induction, SDC increased stepwise following the BDL procedure. Following the recanalization, both the liver ADC and the liver SDC returned to normal values. Data are presented with scatter plot (each dot represents the measure of one rat), median, and 95% confidence interval. ADC unit: ×10^-3^ mm^2^/s; SDC unit: au/s.

**Table 1.**
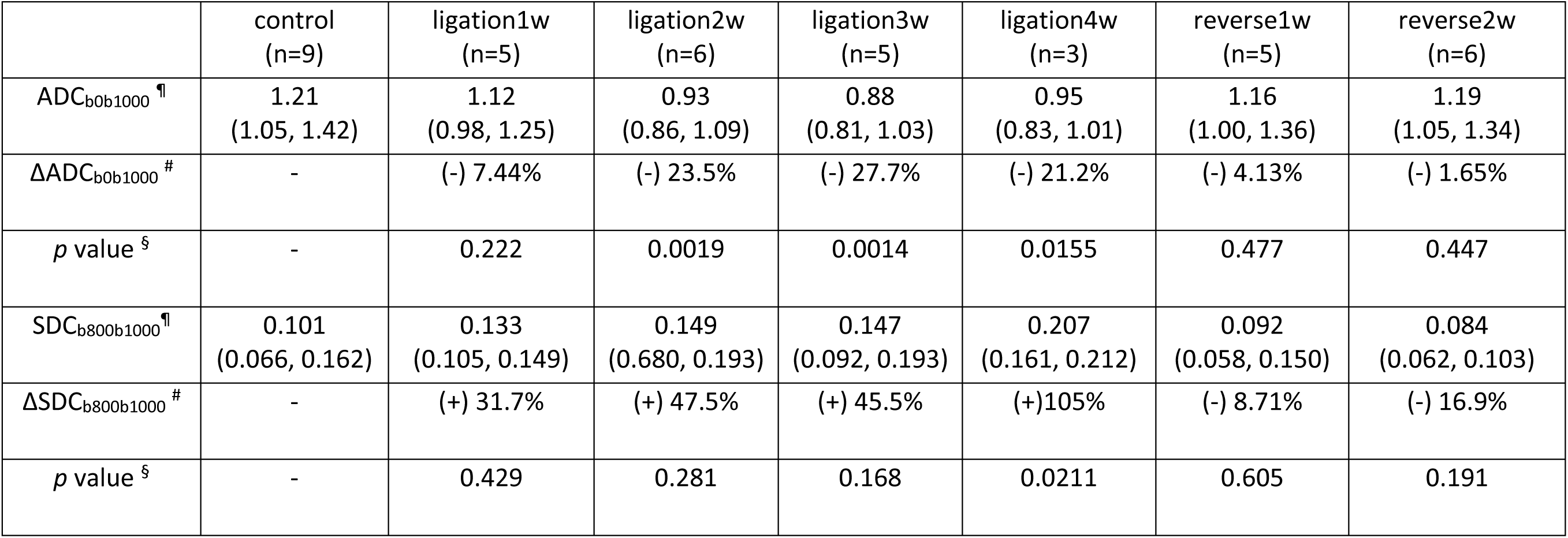
Rat liver ADC (unit: ×10^-3^ mm^2^/s) and SDC (unit: au/s) changes following the BDL procedure and recanalization procedure. ^¶^ : median, 95% CI (confidence interval) ^#^: difference compared to the values of the controls; ^§^ : compared to the values of the controls

|  | control<br>(n=9) | ligation1w<br>(n=5) | ligation2w<br>(n=6) | ligation3w<br>(n=5) | ligation4w<br>(n=3) | reverse1w<br>(n=5) | reverse2w<br>(n=6) |
| --- | --- | --- | --- | --- | --- | --- | --- |
| ADC<sub>b0b1000</sub><sup>¶</sup> | 1.21<br>(1.05, 1.42) | 1.12<br>(0.98, 1.25) | 0.93<br>(0.86, 1.09) | 0.88<br>(0.81, 1.03) | 0.95<br>(0.83, 1.01) | 1.16<br>(1.00, 1.36) | 1.19<br>(1.05, 1.34) |
| ΔADC<sub>b0b1000</sub><sup>#</sup> | - | (-) 7.44% | (-) 23.5% | (-) 27.7% | (-) 21.2% | (-) 4.13% | (-) 1.65% |
| *p* value<sup>§</sup> | - | 0.222 | 0.0019 | 0.0014 | 0.0155 | 0.477 | 0.447 |
| SDC<sub>b800b1000</sub><sup>¶</sup> | 0.101<br>(0.066, 0.162) | 0.133<br>(0.105, 0.149) | 0.149<br>(0.680, 0.193) | 0.147<br>(0.092, 0.193) | 0.207<br>(0.161, 0.212) | 0.092<br>(0.058, 0.150) | 0.084<br>(0.062, 0.103) |
| ΔSDC<sub>b800b1000</sub><sup>#</sup> | - | (+) 31.7% | (+) 47.5% | (+) 45.5% | (+)105% | (-) 8.71% | (-) 16.9% |
| *p* value<sup>§</sup> | - | 0.429 | 0.281 | 0.168 | 0.0211 | 0.605 | 0.191 |

In human dataset 1 and dataset 2, advanced liver fibrosis had an ADC lower than that of early-stage liver fibrosis, on the other hand advanced liver fibrosis had a SDC higher than that of early-stage liver fibrosis (*Figure-2, Figure-3*. *table -2*). In human dataset 3, liver MRI-cirrhosis had an ADC comparable to that of liver fibrosis, whereas liver MRI-cirrhosis had a SDC higher than that of liver fibrosis (*Figure-4*, *table-2*).

**Figure-2.**
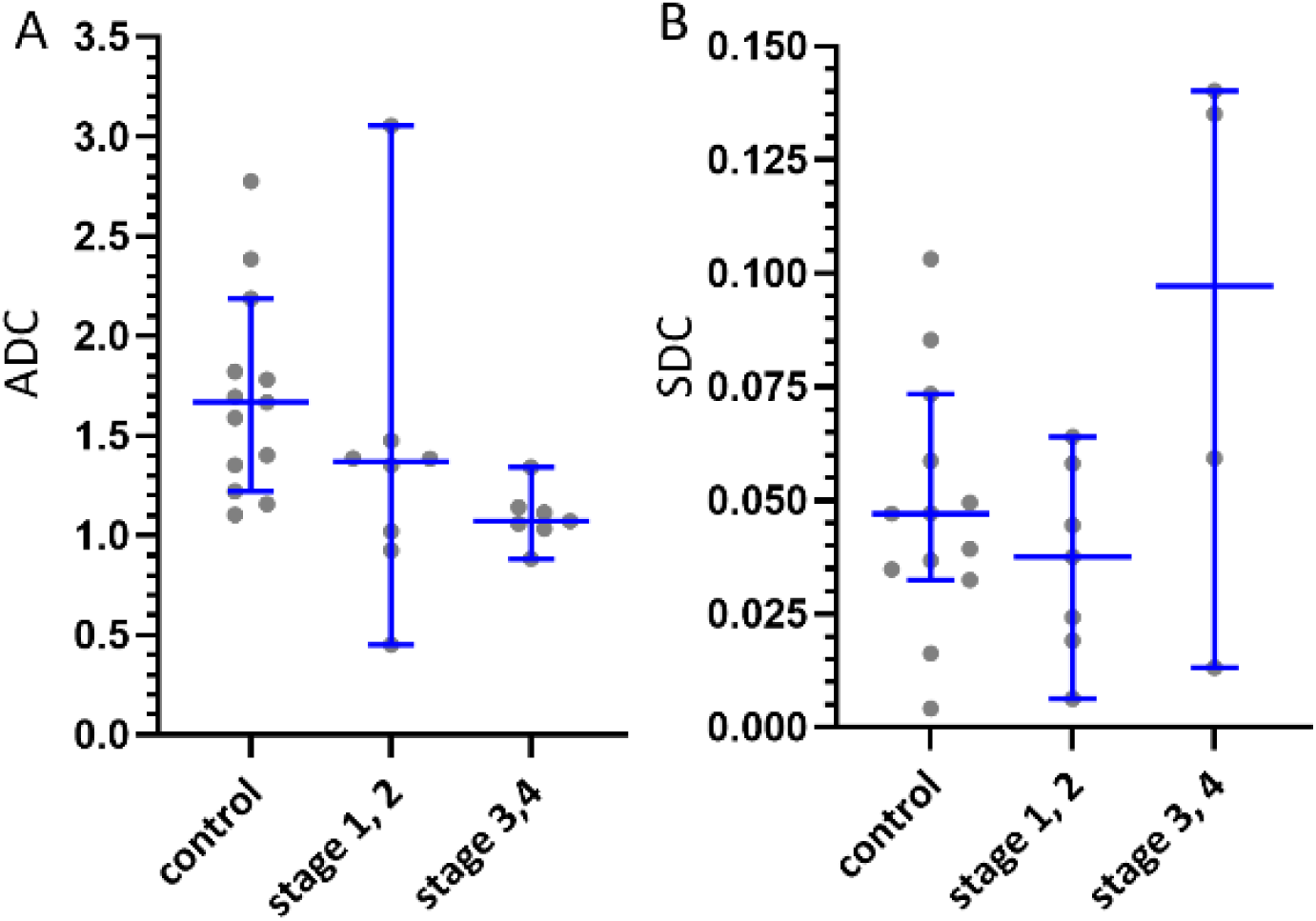
Human dataset 1 liver ADC_b0b600_ and SDC _b400b600_ results. Liver advanced fibrosis (stages 3 and 4) has an ADC lower than that of early-stage liver fibrosis (stages 1 and 2); liver advanced fibrosis has a SDC higher than that of early-stage liver fibrosis. Each dot represents a study subject, and blue lines represent median and 95% confidence interval. ADC unit: ×10^-3^ mm^2^/s; SDC unit: au/s.

**Figure-3.**
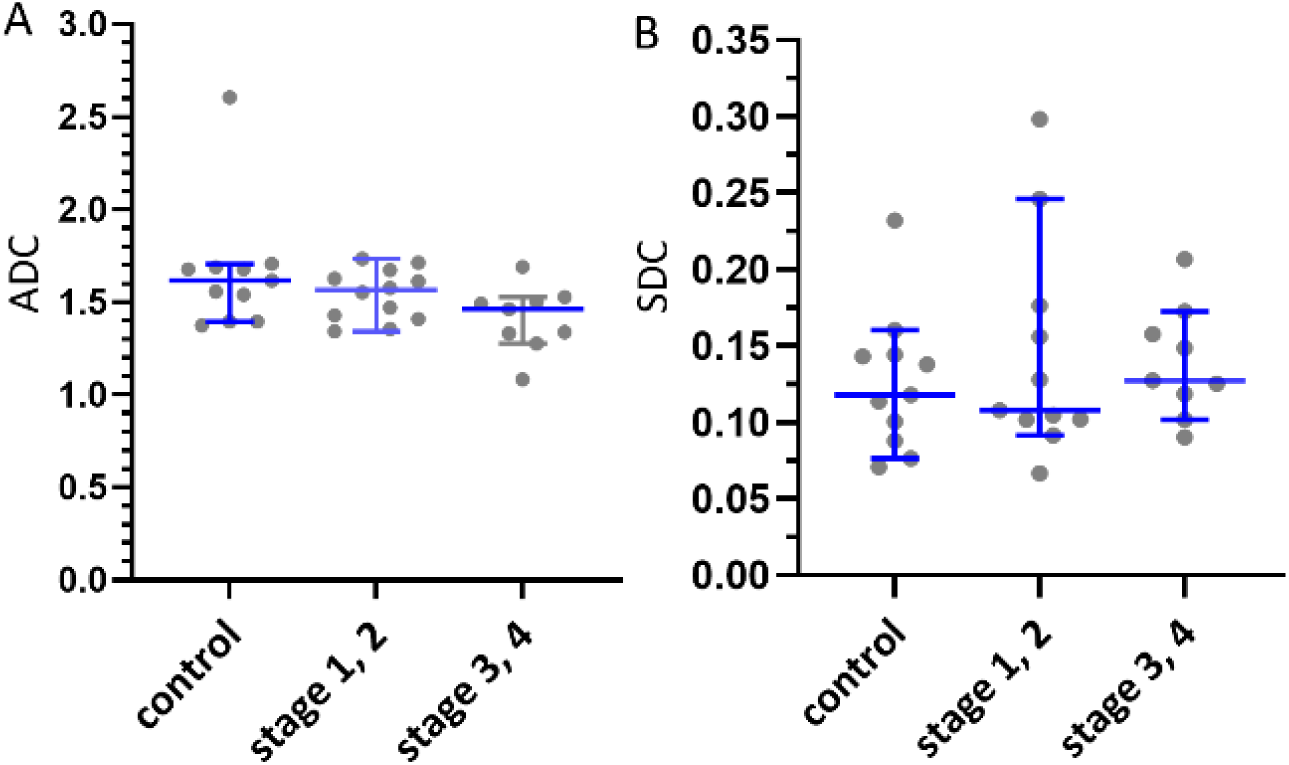
Human dataset 2 liver ADC_b0b600_ and SDC_b400b600_ results. Liver advanced fibrosis (stages 3 and 4) has an ADC lower than that of early-stage liver fibrosis (stages 1 and 2); liver advanced fibrosis has a SDC higher than that of early-stage liver fibrosis. Each dot represents a study subject, and blue lines represent median and 95% confidence interval. ADC unit: ×10^-3^ mm^2^/s; SDC unit: au/s.

**Figure-4.**
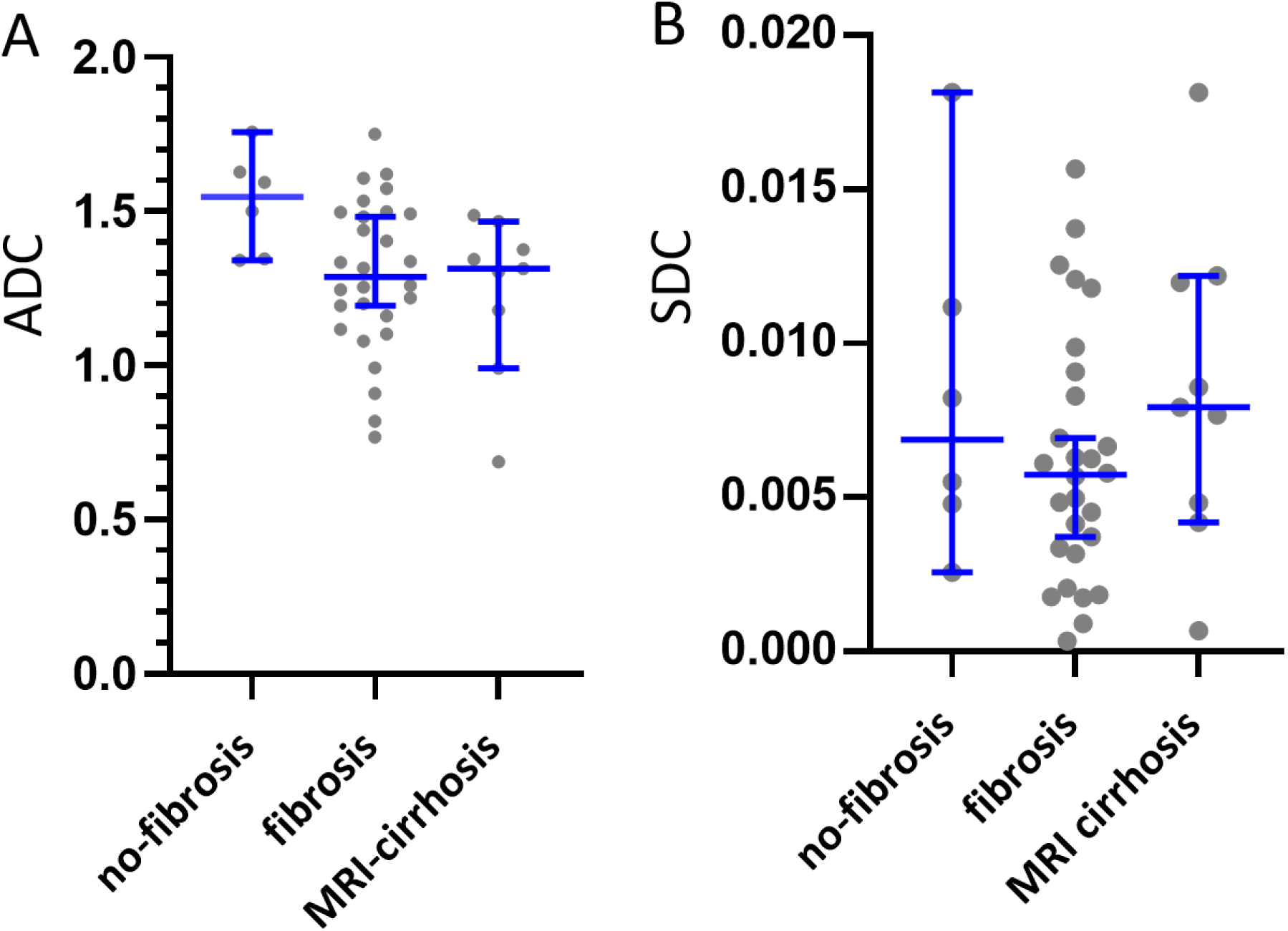
Human dataset 3 liver ADCb0b800 and SDC_b500b800_ results. Livers with MRI-visible cirrhosis had an ADC lower than that of liver fibrosis without MRI-visible cirrhosis. Livers with MRI-visible cirrhosis had an SDC higher than that of liver fibrosis without MRI-visible cirrhosis. Each dot represents a study subject, and blue lines represent median and 95% confidence interval. ADC unit: ×10^-3^ mm^2^/s; SDC unit: au/s.

**Table 2.** Liver ADC (unit: ×10^-3^ mm^2^/s) and SDC (unit: au/s) of three clinical datasets (males only). For datasets 1 and 2, no-fibrosis subjects were healthy controls, and early-stage fibrosis included histological stage 1 and stage 2 fibrosis, advanced fibrosis included histological stage 3 and stage 4 fibrosis. For datasets 3, no-fibrosis subjects were patients with liver focal nodular hyperplasia (FNH), but without fibrosis according to surgical histopathology; all fibrosis patients in datasets 3 had hepatocellular carcinoma (HCC). For datasets 3, advanced fibrosis patients had MRI-visible cirrhosis and early-stage fibrosis did not have MRI-visible cirrhosis. ^¶^ : median, 95% CI (confidence interval). ^#^: Kruskal-Wallis test comparing three groups. ^‡^ : Mann Whitney test comparing subjects without fibrosis and subjects with the early-stage fibrosis. ^§^: Mann Whitney test comparing the early-stage fibrosis subjects and advantaged stage fibrosis subjects. Note that, the absolute values of SDC are highly sensitive to scan parameters and magnet field strength and not directly comparable among these three datasets.

| Data | Parameter | Fibrosis |  |  | p-value |  |
| --- | --- | --- | --- | --- | --- | --- |
|  |  | no-fibrosis | Early-stage fibrosis | Advanced fibrosis |  |  |
| Dataset 1 | ADC<sub>b0b600</sub> ¶ | 1.67<br>(1.22, 2.19) | 1.37<br>(0.45, 3.05) | 1.07<br>(0.88, 1.34) | 0.006 # |  |
|  | SDC<sub>b400b600</sub> ¶ | 0.0471<br>(0.0324 to 0.0374) | 0.0375<br>(0.0063 to 0.0640) | 0.0972<br>(0.0131 to 0.140) | 0.44 ‡ | 0.23 § |
| Dataset 2 | ADC<sub>b0b600</sub> ¶ | 1.62<br>(1.396 to 1.708) | 1.57<br>(1.409 to 1.672) | 1.46<br>(1.28 to 1.53) | 0.067 # |  |
|  | SDC<sub>b400b800</sub> ¶ | 0.118<br>(0.0762 to 0.160) | 0.108<br>(0.092 to 0.246) | 0.110<br>(0.102 to 0.172) | 0.75 ‡ | 0.82 § |
| Dataset3<br>(MRI<br>classification) | ADC<sub>b0b800</sub> ¶ | 1.55<br>(1.34 to 1.76) | 1.30<br>(1.20 to 1.48) | 1.31<br>(0.99 to 1.47) | 0.033 # |  |
|  | SDC<sub>b500b800</sub> ¶ | 0.0069<br>(0.0026 to 0.0181) | 0.0057<br>(0.0037 to 0.0041) | 0.0079<br>(0.0042 to 0.0122) | 0.47 ‡ | 0.24§ |

In all three human datasets, early-stage liver fibrosis had both ADC and SDC lower than those of the controls. If each liver SDC datum was normalized by the mean value of the controls of its respective dataset and three human datasets were summed together, stage-1 liver fibrosis had a normalized SDC value lower than that of the controls, and there was a stepwise increase of SDC value from stage-1 liver fibrosis to stage-4 liver fibrosis (*Figure-5*).

**Figure-5.**
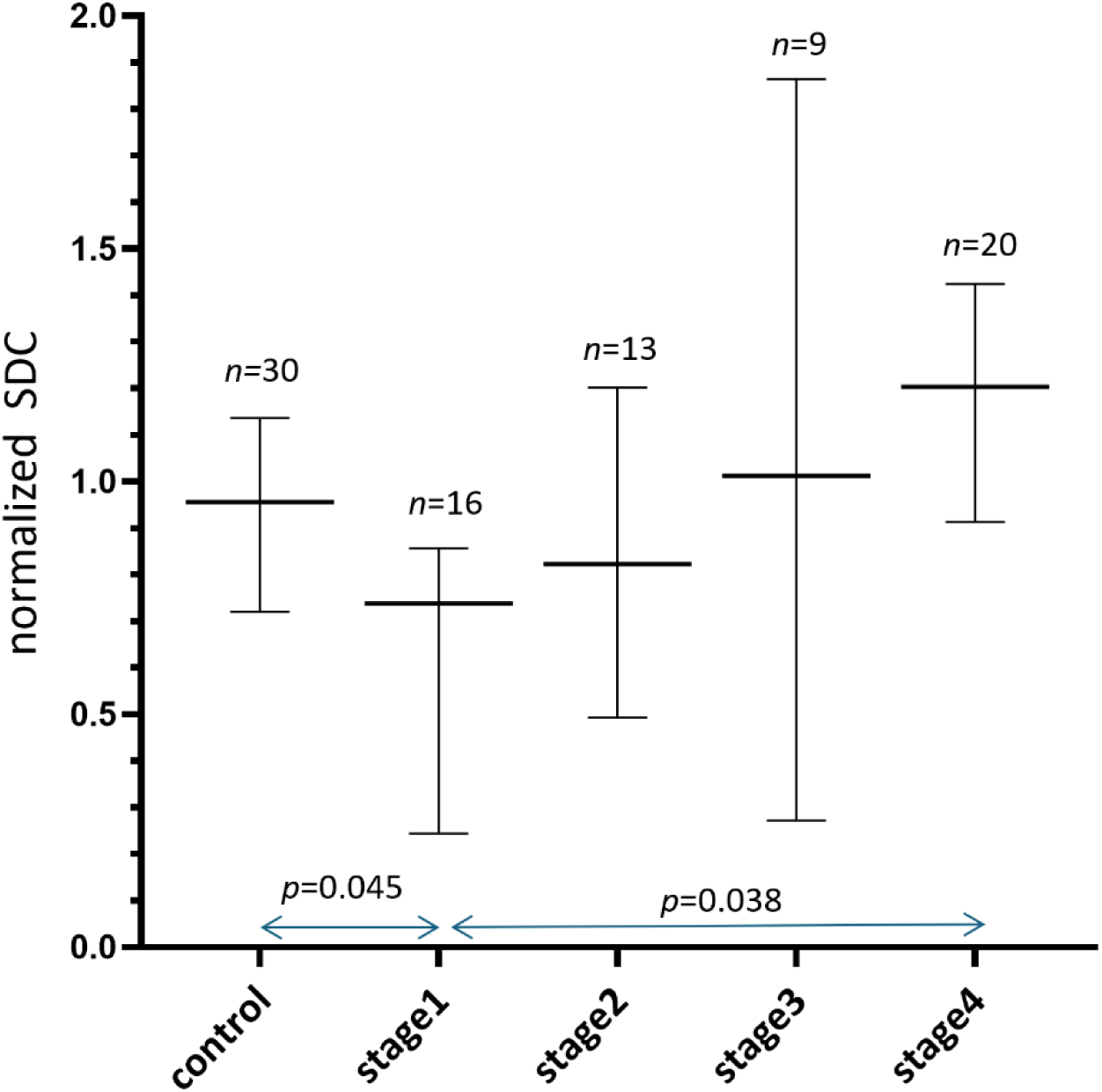
Summed results of normalized SDC values (median and 95% confidence interval) of three clinical datasets. In this graph, all liver SDC values were normalized to the mean value for controls (three datasets normalized separately), and the mean values for controls of each of the three datasets were assumed to be ‘1’. Stage-1 liver fibrosis has a normalized SDC value lower than that of the controls, and there is a stepwise increase in SDC value from stage-1 liver fibrosis to stage-4 liver fibrosis (i.e., cirrhosis).

## Discussion

ADC measure is affected by the tissue T2 [13–16]. Eq. (1) for ADC calculation shows, the greater the signal loss between S(*b_1_*) and S(*b_2_*), the higher the ADC value. Although the low *b*-value can be nominally 0, in reality, the gradient switching for spatial encoding in the slice selection and readout directions always generates *b*-values greater than 0. This makes it impossible to acquire images with exactly *b*=0 s/mm^2^. The application of diffusion gradients leads to the tissue demonstrating shorter ‘apparent’ T2 relative to the T2 measured when *b*=0 s/mm^2^. For moving spins, the application of the pair of diffusion gradients will contribute to residual dispersion of ‘spin focusing’ in the transverse plane (even after the application of the second ‘re-focusing’ diffusion gradient), leading to a shortening of the measured T2 [16, 17]. It is reasonable to assume that, at the voxel level, there will be ‘ever-existing’ moving spins, which are also contributed by irregular macroscopic body motion. When the motion probing gradient *b*-value is increasingly high, even for assumed static voxels, the ‘re-focusing’ by the second diffusion gradient is increasingly less coherent [16]. It has been shown that, if we use Eq. [1] and apply a high *b*-value and a higher *b*-value images (such as 400 and 600 s/mm^2^) to calculate ADC, the ‘T2 shine-through’ of ADC could not be resolved [19]. The IVIM parameter D_slow_ also suffers from ‘T2 shine-through’ effect [15]. For example, the same as ADC, IVIM-D_slow_ shows a lower value in HCC compared to adjacent liver tissues and a lower value in fibrotic livers compared to normal livers [19, 26, 27]. IVIM-D_slow_ of the spleen has been consistently measured much lower than that of the liver [28]. SDC was proposed to mitigate the difficulties associated with the ‘T2 shine-through’ effect of ADC and IVIM-D_slow_. SDC is negatively associated with ADC for tissues with T2 shorter than 70 ms, positively associated with ADC for tissues with T2 longer 70 ms. The strength of the correlation will depend on the T2 of the tissue [15, 16]. It was shown that, an increase in TE for DWI acquisition led to an increase in liver ADC, whereas an increase in TE for DWI image acquisition led to a decrease in liver SDC. Moreover, the magnitude of changes (increase or decrease) was much greater for the liver than for the spleen with liver having a much shorter T2 than the spleen (liver: 40 ms, spleen: 60 ms, 3.0T) [23, 29]. Note that, from MRI signal decay point of view, a longer TE is equivalent to a shorter T2. For brain gliomas and parotid gland tumors (both have a mean T2 > 70ms), a positive correlation of Pearson *r* of around 0.6 ∼ 0.7 have been noted [30, 31].

SDC has shown practical advantage in various scenarios. For example, in a study of 63 patients with diffuse gliomas [30 IDH (Isocitrate dehydrogenase)-mutant and 33 IDH-wildtype], SDC_b500b750_ separated IDH mutant negative tumors and mutant positive tumors with an AUROC (area under receiver operating characteristic curve) of 0.828. ADC_b0b1000_ separated IDH mutant negative tumors and mutant positive tumors with an AUROC of 0.760 for separation [32]. Thus, SDC as a biomarker offers a better differentiation power than ADC. A combination of diffusion-derived ‘vessel density’ (DDVD) [20, 33], ADC, and SDC achieved an AUROC of 0.9 for separating IDH-mutant and IDH-wildtype gliomas [30]. In studies of parotid gland tumors, DDVDr, SDCr, and ADCr were the metrics of the tumor divided by the metrics of tumor free parotid gland tissue. A combination of ADCr_b0b800_, SDCr_b600b800_, and DDVDr_b20_ separated parotid gland malignant tumors and benign tumors (i.e., Warthin’s tumors and pleomorphic adenomas) with an AUROC of 0. 805 [31]. Based on the T2 weighted image signal and these three metrics of DDVD, SDC, and ADC, we applied a scoring scheme termed ‘LiverMss-FNH’ to evaluate liver solid mass. In two studies totaling 25 focal nodular hyperplasia (FNH) and 132 liver malignant tumors, LiverMss ≥ 3.0 suggests the possibility of a liver mass being FNH, and LiverMss ≥ 4 can strongly favor the diagnosis for FNH [22, 34]. A typical SDC signal, i.e., being iso-signal or slightly high signal (but not high signal), has an odd ratio of 38 in favor of liver FNH over liver malignant tumors [22, 34]. More recently, we showed that a liver mass with an iso-signal or slightly high DDVD signal while with an SDC higher or equal to that of the kidneys had an odds ratio of 34.7 in favor of intrahepatic cholangiocarcinoma over HCC [35], and a higher SDC value is associated with a better intrahepatic cholangiocarcinoma patient survival potential [36].

The lowered value of ADC liver fibrotic livers is likely due to at least three factors, i.e., 1) longer T2 of fibrotic livers is associated a less signal decay between S(*b_1_*) and S(*b_2_*) according to Eq [1]; 2) increased iron-related susceptibility depresses ADC measure [37, 38]; and 3) liver fibrosis/cirrhosis are associated decrease blood perfusion to the liver [20, 33, 39, 40]. In studies observing the aging effect on ADC and SDC, we noted that decreased blood perfusion to the liver in the older subjects leads to lower liver ADC but has no effect on SDC [23, 41]. In the current study, the rat BDL model results show liver ADC decreased stepwise following the duration (in weeks) of BDL, whereas SDC increased stepwise. In all human data, advanced liver fibrosis had an ADC lower than that of liver early-stage fibrosis patients, on the other hand advanced liver fibrosis had a SDC higher than that of liver early-stage fibrosis. The latter is consistent with the clinical observation that more severe liver fibrosis is associated with higher liver water level (particularly ECW), and this in turn would relate to faster tissue diffusion. It is noted that early-stage fibrosis has a SDC lower than that of the healthy controls. We attribute this to the fact that liver fibrosis is associated with increased liver iron level [42]. SDC measure appears to be highly sensitive to iron/susceptibility effect. For example, compared to pre-menopausal women, post-menopausal women after 50 years have an increase in liver iron level and a lowering of liver SDC, whereas this change is not observed in similarly aged men [23]. It was well established that regardless of the aetiology, iron-loading is frequently observed in chronic liver diseases. Increased liver iron in viral hepatitis may be a combined consequence of dysregulated liver iron homeostasis and normal defensive processes adopted during infections which involves sequestration of iron by hepatic cells to limit access to pathogens to inhibit their proliferation [43]. Excess iron can induce fibrosis-promoting signals in the parenchymal and non-parenchymal cells which accelerate disease progression and exacerbate liver pathology [44, 45]. Normal liver iron concentration is lower than 35 μmol/g of dry weight [46]. When liver iron concentration crosses a threshold of 60 µmol/g, hepatic stellate cells’ functionality begins to derail, and when it exceeds 250 µmol/g, cirrhosis becomes inevitable [46]. Despite the iron-related susceptibility is likely to be more apparent for higher grade fibrosis, the effect of elevated water content might surpass that of the susceptibility effect for cirrhosis or advanced stage fibrosis, finally leading to an elevated SDC (*Figure-6*).

**Figure-6.**
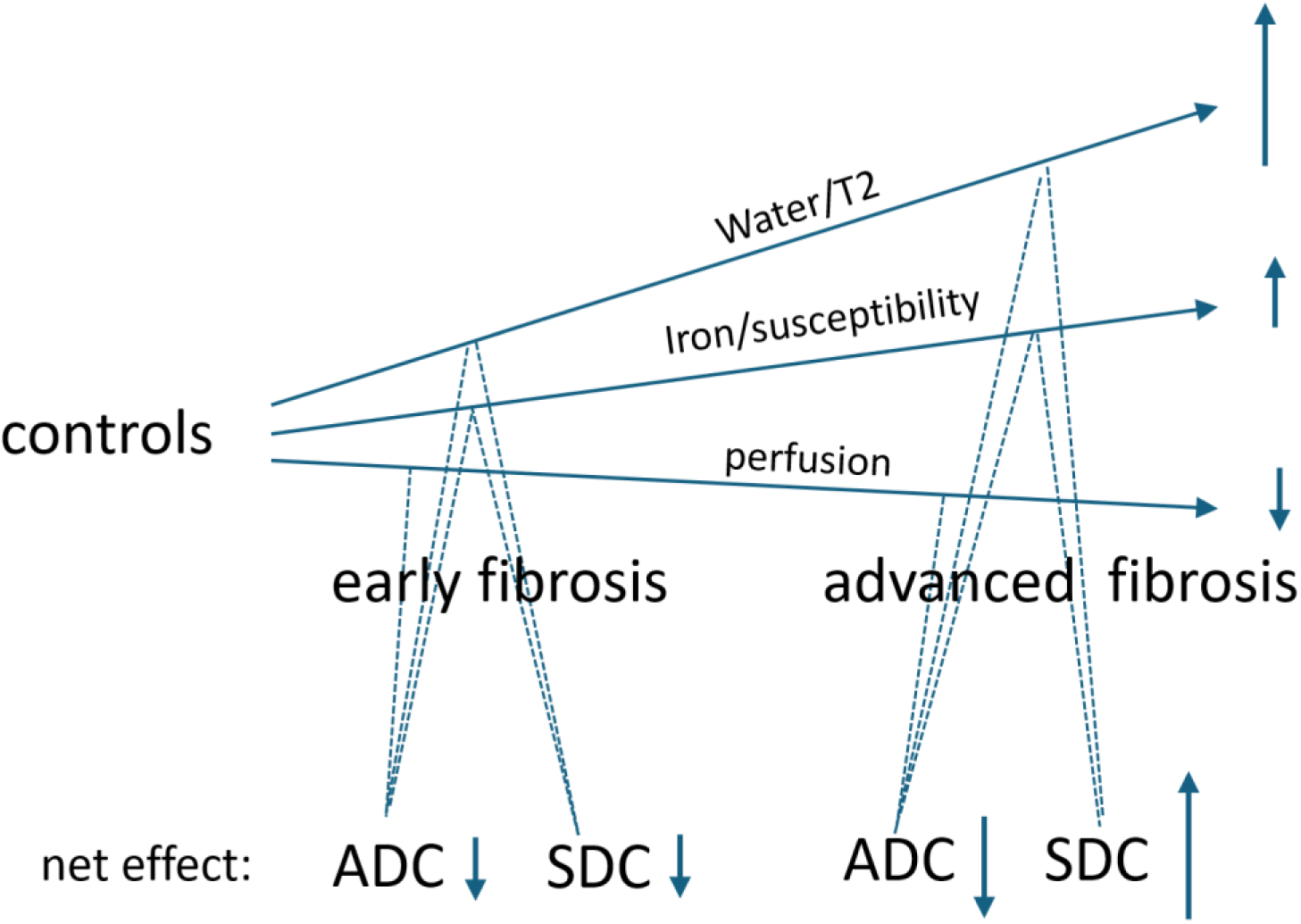
Postulated impact of three factors of perfusion, iron/susceptibility, and water content on ADC and SDC. Liver fibrosis is associated with lower perfusion [20, 33, 39, 40], higher iron/susceptibility [42–46], and higher water content [1–7]. These three factors all contribute to lower ADC measure, and more so with advanced fibrosis. Perfusion generally does not affect SDC [23, 41]. Higher iron/susceptibility lowers SDC measure [23], whereas higher water content elevates SDC measure. For early-stage liver fibrosis, the net effect of susceptibility and water leads to a lower SDC; while for advanced liver fibrosis, the net effect of susceptibility and water leads to a higher SDC.

There are many limitations to this study. This study conveniently re-used three sets of our historical liver IVIM data, while the liver iron level and its effect magnitude on SDC were not quantified and could only be postulated. However, it is reasonable to assume the liver iron level in our human data is: fibrotic liver > normal liver [40–44]. The iron/susceptibility effect may not be elevated in the BDL animal model. Water content in the liver was also not quantified for the liver in this study. Literature has well documented that advanced stage liver fibrosis has longer T2 and higher ECW/TBW [5–7]. The sample size is limited for all the four datasets analysed. Despite statistical significance not being achieved for individual human datasets, the trend was the same for all three datasets, and statistical significance was achieved when we combined the results of the three datasets together (*Figure-5*). Finally, because a higher proportion of liver fibrosis/cirrhosis was in their early 50’s, we only used males’ clinical data in this study, to tease out the potential menopausal effect on liver SDC in females’ clinical data will require another study.

In conclusion, rat liver fibrosis model shows liver ADC decreases following BDL surgery, whereas liver SDC increases. Three sets of human data consistently show that advanced liver fibrosis has a SDC higher than that of early-stage liver fibrosis. These SDC results are consistent with the known pathophysiology that patients with cirrhosis have sodium/water retention, and liver fibrosis/cirrhosis is associated with a higher liver water content and longer T2.

## list of abbreviations

ADC: apparent diffusion coefficient
BDL: biliary duct ligation
DDVD: diffusion-derived vessel-density
DEN: diethylnitrosamine
DWI: diffusion weighted imaging
ECW: extracellular water
FOV: field of view
FNH: focal nodular hyperplasia
HCC: hepatocellular carcinoma
IDH: Isocitrate dehydrogenase
IVIM: Intravoxel incoherent motion
NEX: Number of excitations
ROIs: regions-of-interest
SDC: slow diffusion coefficient
SPIR: Spectral Pre-saturation with Inversion-Recovery
TR: time of repetition
TE: time of echo
TBW: total body water

